# Genomic insights into the domestication of the chocolate tree, *Theobroma cacao* L

**DOI:** 10.1101/223438

**Authors:** Omar E. Cornejo, Muh-Ching Yee, Victor Dominguez, Mary Andrews, Alexandra Sockell, Erika Strandberg, Donald Livingstone, Conrad Stack, Alberto Romero, Pathmanathan Umaharan, Stefan Royaert, Nilesh R. Tawari, Ng Pauline, Ray Schnell, Wilbert Phillips, Keithanne Mockaitis, Carlos D. Bustamante, Juan C. Motamayor

**Author notes:** Corresponding authors: Carlos D. Bustamante and Juan Carlos Motamayor.

## Abstract

Domestication has had a strong impact on the development of modern societies. We sequenced 200 genomes of the chocolate plant *Theobroma cacao* L. to show for the first time that a single population underwent strong domestication approximately 3,600 years (95% CI: 2481 – 10,903 years ago) ago, the Criollo population. We also show that during the process of domestication, there was strong selection for genes involved in the metabolism of the colored protectants anthocyanins and the stimulant theobromine, as well as disease resistance genes. Our analyses show that domesticated populations of *T. cacao* (Criollo) maintain a higher proportion of high frequency deleterious mutations. We also show for the first time the negative consequences the increase accumulation of deleterious mutations during domestication on the fitness of individuals (significant negative correlation between Criollo ancestry and Kg of beans per hectare per year, P = 0.000425).

---

Organized state societies were only possible after the development of agriculture, which involved the domestication of numerous plants and animals^1, 2^. We are just starting to understand this process more from a genetic perspective thanks to expanding genomic technologies. Of particular interest is identifying the demographic scenario, timeline for domestication and genomic consequences for species that have been critical in the development of societies^2–4^ Among all species, there is a special place in the general culture for the domestication of *Theobroma cacao* L, the plant from which chocolate is made. The chocolate tree has played a fundamental role in the development of Mesoamerican civilizations^5^ and has been a topic of research for over 100 years, but its domestication history has remained a controversial topic ^6–8^. While the domestication history of cacao has sparked the interest across diverse disciplines, our knowledge of the process is incomplete and often involves partial information focused on either a few genetic groups, a few geographical regions, fragmented archeological information, or a limited number of genetic markers^8–12^. In this work we report and analyze whole genome variation of 200 *T. cacao* individuals (Supplementary Table 1) to investigate the evolutionary origin of Criollo, the cacao tree domesticated in Mesoamerica. We also examine the consequences of the domestication process in the genomic architecture of the accumulation of deleterious mutations along the genome, which in turn allowed us to understand critical limits of Criollo (domesticated) cacao productivity.

Current dogma suggests cacao was introduced to Mesoamerica in Olmec times from cacao varieties present in the Upper Amazon (Northern South America), the hotbed of diversity for the species^6, 8^. Anthropological research, in particular, supports this view ^6, 9, 12^. In addition, the continuous intermixing of farmed and wild cacao trees has likely continued to shape both gene pools in recent times ^11, 13, 14^ Both, the impact of ancient domestication processes and modern hybridization on the genetic variation in the species is largely unknown in *T. cacao.*

Re-sequencing of 200 accessions at high coverage (average coverage 22X) generated ~4.52 Trillion base pairs. After aligning reads to the cacao reference (Matina-v1.1^15^), we identified 7,412,507 single nucleotide polymorphisms (SNPs). We find that cacao presents a high genetic variability of ~5 SNPs per kilobase per individual, which is similar to what has been observed in Arabidopsis^16, 17^. Although the large majority of identified variants are noncoding, we identified 322,275 missense variants and 220,043 synonymous variants in 29,408 genes. We also identified 10,062 variants predicted to change the splice donor sites, which could be responsible for polymorphism impacting the number of transcripts produced ^15^. Among the potential changes that alter the length of the transcripts, we identified 8,470 start losses, 19,956 stop gains, and 8,588 stop losses. Overall, this SNP dataset represents a new resource for cacao biology that we hope will accelerate breeding programs (for a catalog of SNPs contextualized with respect to gene annotation see supplementary table 2 and figure S1).

Model-based clustering analysis enabled us to identify 10 genetic clusters, consistent with previous analyses^10^ and allowed us to correctly assign overall ancestry to previously uncharacterized accessions (Figure 1A). We also present, for the first time, the characterization of the global ancestry assignments for the hybrids revealed the relative contribution of these 10 groups to natural admixed individuals and man-made hybrids (Figure 1A). Our results show that there is an overrepresentation of genetic material from Amelonado, Criollo and Nacional ancestry in the majority of admixed individuals. There is a concomitant underutilization of other genetic groups in current agronomist practice, which provides “blue ocean” opportunities for crop improvement (see supplementary information).

**Figure 1.**
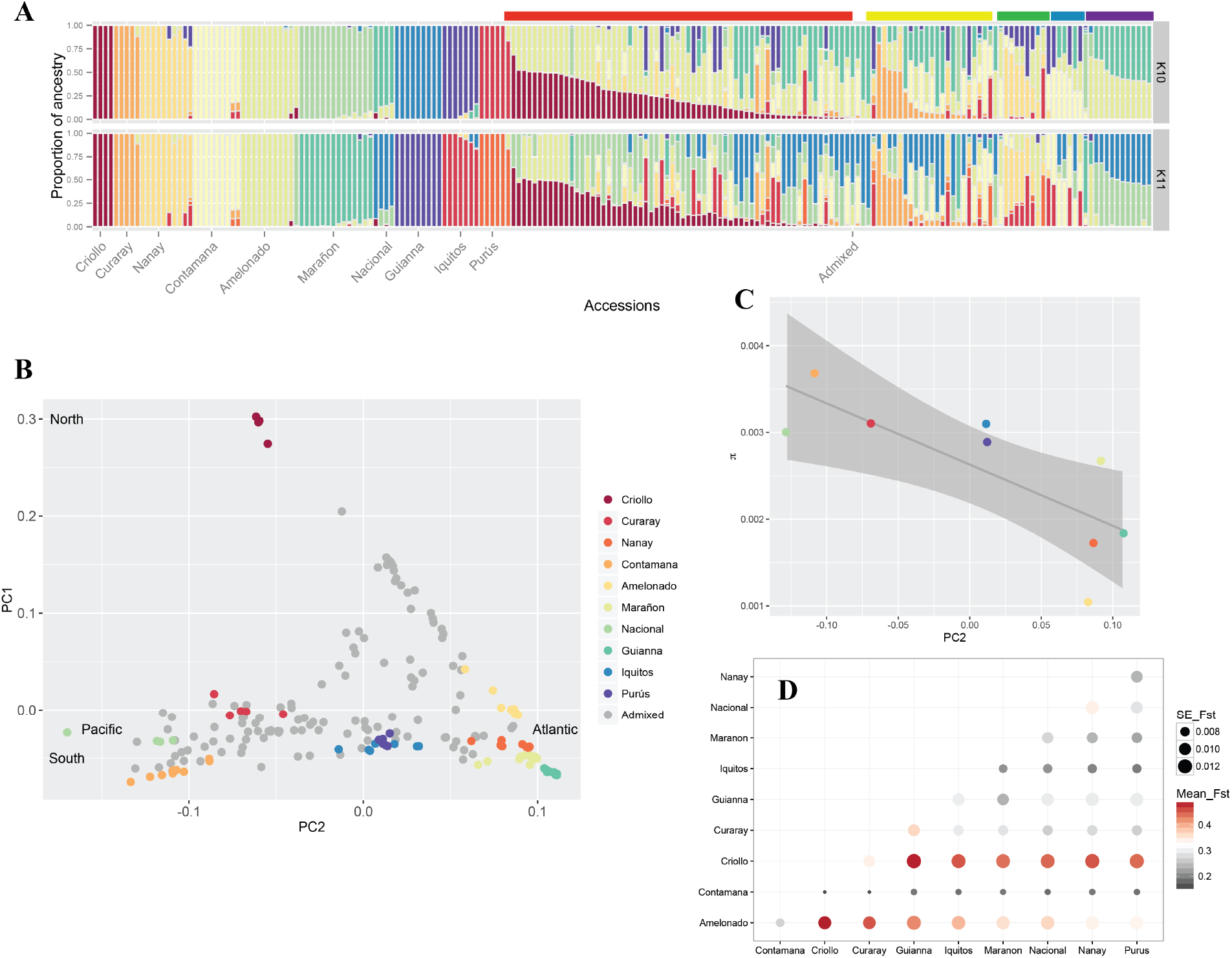
Population Genetic structure in *T. cacao.* **A**, The 10 main genetic clusters can be recovered (top panel) although further structure (11 clusters) seems to be meaningful as a considerable number of admixed individuals present ancestry from a subset of Amelonado ancestry (bottom panel). Color bars on top of the admixed individuals show our suggested grouping for the hybrids. **B**, MDS showing a gradient of differentiation form the West to the East side of the Amazon (PC2) and a major separation of the Criollo group that corresponds to the Mesoamerican domesticated group (PC1). **C**, Significant decay of genetic diversity 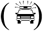 for the species along PC2 is supportive of the origin of the species being in the western side of the Amazon basin (Criollo is excluded). **D**, All 10 population genetic groups that have been described for the species are highly differentiated, with Criollo presenting a larger average Fst when compared against all the other groups.

Further analysis of the population structure shows that the Criollo group is clearly differentiated from the rest of the genetic clusters along the first axis of a multi-dimensional scaling (MDS) analysis (Figure 1B). The second component of the MDS analysis presents a gradient separating genetic clusters roughly from Pacific to Atlantic (bottom to top of the second component), consistent with a natural process of differentiation from the higher diversity groups on the Pacific side of the Amazonian basin to those of lower diversity on the Atlantic side (See Figure 1C, excluding domesticated Criollo, Y = 0.202 – 72.71X, p=0.02). It has been proposed that the center of origin of the species is located in the western Amazon^7, 18^. Our observation of the significant decline in genetic diversity from Pacific to Atlantic is consistent with the suggested center of origin for the species. The spread of admixed individuals in the MDS space is consistent with our admixture analysis in which individuals fall into two general categories: i) those that present admixture between Criollo and Amelonado with a minor contribution from other groups; and ii) those hybrids that present admixture along the Atlantic-Pacific gradient. There is a pattern of strong differentiation between all genetic clusters (Figure 1D, F_ST_ values range between 0.16 and 0.65), with a larger differentiation between Criollo and any other group, consistent with either a scenario of strong drift during a recent process of domestication or an old diversification of the Criollo population from the rest of the genetic clusters. Given previous anthropological and genetic evidence, it is more likely that this pattern is the result of strong domestication from a small pool of seeds that were used to create the Criollo group.

The hypothesis of a single event of domestication (along with genetic drift after transport from South to Central America) predicts that Criollo would show a higher differentiation to other groups than that observed between any other pairwise comparison of the populations. Our population structure analyses are consistent with this prediction (Figure 1B, D). Our model-based analysis of population differentiation provides evidence that Criollo was the result of a single domestication event, undergoing extreme drift after separating from its most closely related population (represented as a longer branch in Figure 2A,B). This analysis also shows that Criollo is most closely related to Curaray suggesting that the origin of domesticated cacao was a subset of ancient Curaray germplasm ^10^, a genetic cluster that has been described for the North of Ecuador and South of Colombia ^10, 19^. After exploring multiple models of differentiation with admixture, we found no evidence supporting subsequent contributions of any group to the domesticated Criollo (Figure 2B). However, we learned from this analysis that multiple instances of admixture have potentially occurred among multiple groups during their natural process of differentiation along the Amazon Basin (see supplementary material).

**Figure 2.**
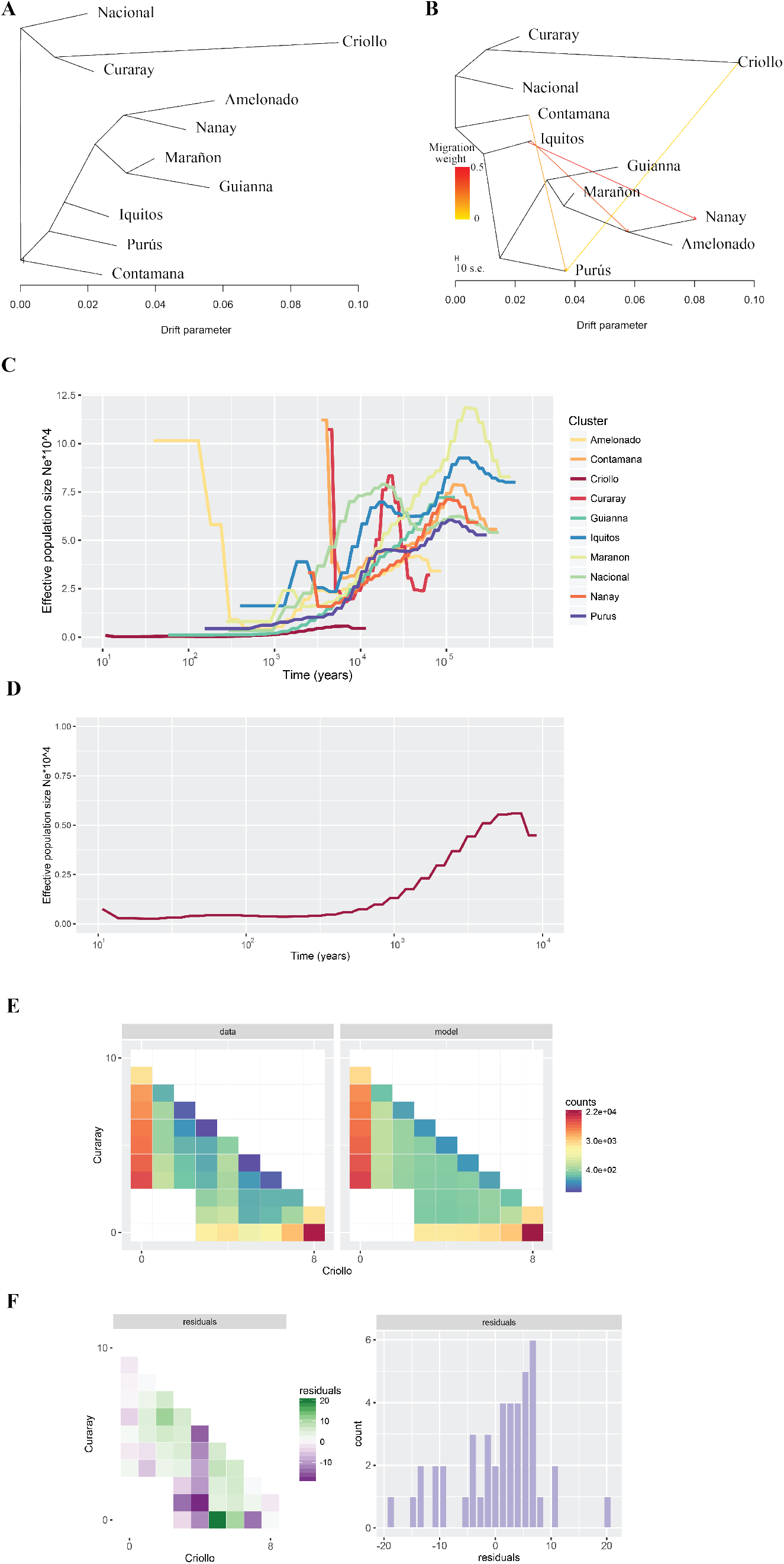
Population Demographics of *T. cacao.* **A**, Maximum likelihood tree generated by *TreeMix* using intergenic regions of whole-genome sequencing data from individuals belonging to each one of the 10 main genetic groups. **B**, Maximum likelihood tree allowing for admixture, as generated by *TreeMix,* showing some of the most significant ancestral contributions (migrations) from and to other groups. **C**, Changes to effective population sizes over time, inferred under the coalescent with PSMC, for each on the 10 genetic groups in cacao. Each line represents the within-population median estimate, smoothed by fitting a cubic spline. **D**, detail of PSMC effective population size reconstruction for Criollo cacao, represented at a different scale to better represent the population decline. **E**, Observed two-dimensional site frequency spectrum (sfs, left panel) for the Criollo/Curaray population pair and expected sfs (right panel) under the inferred demographic model depicted in **F**. The colors correspond to magnitudes (number of SNPs in each minor allele frequency bins). In the bottom panels the Anscombe residuals (different between observed and expected) per frequency bin and as an overall distribution.

In addition to admixture and population differentiation analyses, we investigated the demographic history of the 10 genetic clusters to understand the natural demographic process that has characterized the species historically. Given the relatively small number of accessions per group, we performed analysis with pairwise sequentially Markovian coalescent (psmc) model^20^ which allows evolutionary history inference by analyzing single diploid genomes. In general, the evolutionary history of *T. cacao* shows a common trend towards the reduction of population size/genetic diversity with time (Figure 2C,D). The median demographic history among accessions was used to show the overall trends for the evolutionary history of the groups. The process of reduction of effective population size started prior to the peopling of the Americas, which suggests that the overall reduction of genetic diversity in the species could be tied to environmental changes or historical changes in the distribution of pollinators and/or animals that are involved in the dispersion of the seeds during the Last Glacial Maximum. This result is consistent with recent studies suggesting that most groups of *Theobroma cacao* could have diversified during the last glaciation ^19^. The fact that populations of cacao have been declining historically is consistent with recent studies that have analyzed the conservation status of more than 15,000 species of Amazonian trees, predicting that *T. cacao* could suffer an additional 50% population decline in the near future ^21^.

Using what we learned from the model-based analysis of differentiation (Figure 2A,B) and our overall assessment of the demographic history of the populations (Figure 2C,D) we explored the evolutionary history of domestication of the Criollo group from a Curaray ancestor to answer two critical questions: first, how long ago did the ancestral Curaray populations give rise to what is today known as the Criollo group? and second, how small was the founding population, of Curaray ancestry, used to domesticate cacao Criollo in Central America? For this, we analyzed the frequency spectrum of variants under a model of isolation with migration under a maximum likelihood framework^22^. Our analyses show that the fraction of the ancestral Curaray effective population used to domesticate Criollo in Mesoamerica was indeed very small and comprised of approximately 738 individuals ( 95% CI: 437 – 2,647 individuals). More importantly, we provide for the first time, strong support from genomic data analysis that this process started 3600 years BP (95% CI: 2481 – 10,903 years BP). The observed distribution of shared variants for different minor allele frequency categories (Figure 2E) fits well the predicted values under the best fitted model (Figure 2F); with a reasonable distribution of residuals. Our estimates for the time of separation of the Curaray and Criollo groups overlaps well with archeological evidence and what has been thought to be the onset of cultivation of Criollo in Mesoamerica^8,9,12,23^. These results are consistent with findings of theobromine in Olmec pottery from the capital San Lorenzo as old as the Early Preclassic (1800 – 1600 BCE) ^9, 24^ Our demographic analyses are also consistent with large scale analyses of modern and ancient DNA which pinpoint the colonization of the American continent by humans to roughly 13,000 years ago^25–27^. In addition, recent analysis of the post-colonization demography of human in South America is consistent with human populations staying in relative low numbers for the first 8000 years and then, with the advent of agriculture and thus sedentism, finally experiencing a population expansion at ~ 5Kya similar to that experienced during the Neolithic revolution elsewhere on the globe^28^. In short, our understanding of human demographic history suggests that our inference of *T. cacao* domestication in Mesoamerica between 2481 and 10,903 years BP are strongly consistent with the history of human settling in the region.

One of the greatly appreciated features of the domesticated Criollo cacao is the white cotyledon of the bean that seems to be associated with desirable flavor qualities. Early work has suggested that decreased concentrations of polyphenols, methylxanthines and anthocyanin precursors concentration in the cotyledon are associated with this observation^29–31^. Polyphenols and methylxanthines are responsible for the astringency and bitternes detected in cacao beans^32^, and it is thought that the modification of these compounds during the process of fermentation contributes to the final flavor of chocolate^32^. In fact, during the process of fermentation, the concentration of polyphenols is reduced by up to 70%^33^. Plants of the Criollo variety were likely selected during domestication to reduce this bitterness. We investigated the impact of artificial selection during domestication on the Criollo genome by looking for regions of increased differentiation between Criollo and its sister population Curaray, using methods based on the comparison of the distribution of allelic variation along the chromosomes^34^. We found several regions of the genome in which natural selection has produced higher differentiation between Curaray and Criollo than expected by demographics alone (Figure 3). The most interesting result derives from the identification of genes encoding Laccase 14, Laccase/Diphenol oxidase.

**Figure 3.**
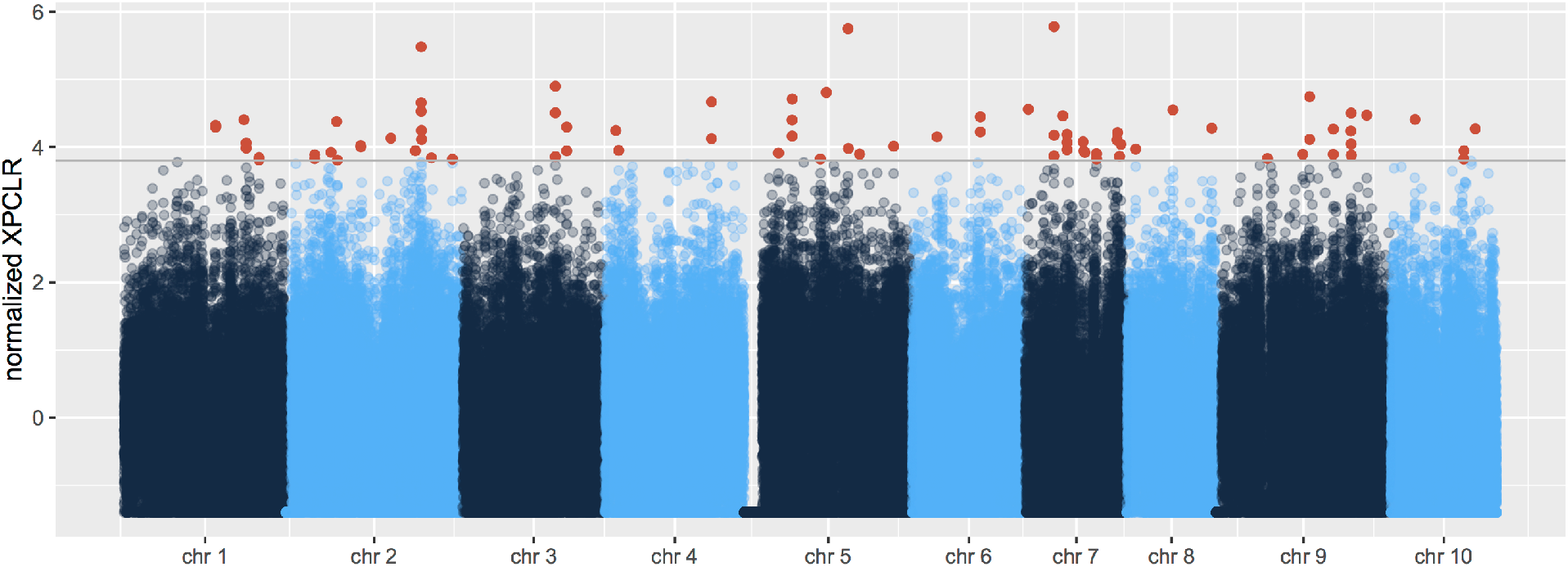
Evidence of positive selection in domesticated *T. cacao.* **A**, Maximum likelihood approach for detecting regions of the genome that diverged significantly from the demographic depicted by the site frequency spectrum in Figure 2E. Red points correspond to windows putatively under selection.

Laccases are normally associated with the process of lignification, but it has recently shown that laccasses are also involved in the metabolism of polyphenols and we hypothesize that selection on these genes likely results in the reduction of the concentration of polyphenols in cacao. We also identify signatures of selection in a region containing the Xanthine dehydrogenase 1 gene, likely involved in the metabolism of methylxanthines (like theobromine) and also likely to have been the result of the process of selection for reduced bitterness^31^. An additional list of genes in regions identified under selection is provided in the supplementary materials (table S3) and include functions involved in genomic stability (Structural maintenance of chromosomes), disease resistance and abiotic stress response (WRKY DNA-binding protein), transcriptional regulation (MYB domain), and signaling (cysteine-rich RLK receptor as well as S-domain-2 5 genes).

Most mutations that appear in the genome are deleterious and have the potential for reducing reproductive success^35–37^. The fate of these mutations and their transit time in a population strongly depends on the intensity of genetic drift, purifying selection, and the degree of dominance of the mutations. Mathematical population geneticists were long concerned with the impact of the accumulation of deleterious mutations in a population^38^. The process of domestication in animals and plants have been used as a framework to study how intense selection of some desirable traits affects the accumulation of deleterious mutations in the population^3, 39^. Yet, thus far, we have little evidence of how the process of accumulation of deleterious mutation affects traits associated with fitness or, in the case of crops, productivity.

Populations of cacao have been declining over time and a natural consequence of the reduction in population size is an increase in inbreeding. Because all 10 populations of cacao are experiencing reductions in population size, it is expected that this process will have a similar effect across populations, and the differences in magnitude of inbreeding will reflect differences in population size. We observe an increase in the amount of inbreeding (estimated as F statistics ^40^) when the admixed cluster of individuals (expected to have low inbreeding) is compared to the naturally defined genetic groups (Figure 4A). These differences between the coefficients of inbreeding can be partially explained as a function of the differences in historical population size among genetic clusters (Figure 4B, see supplementary text).

**Figure 4.**
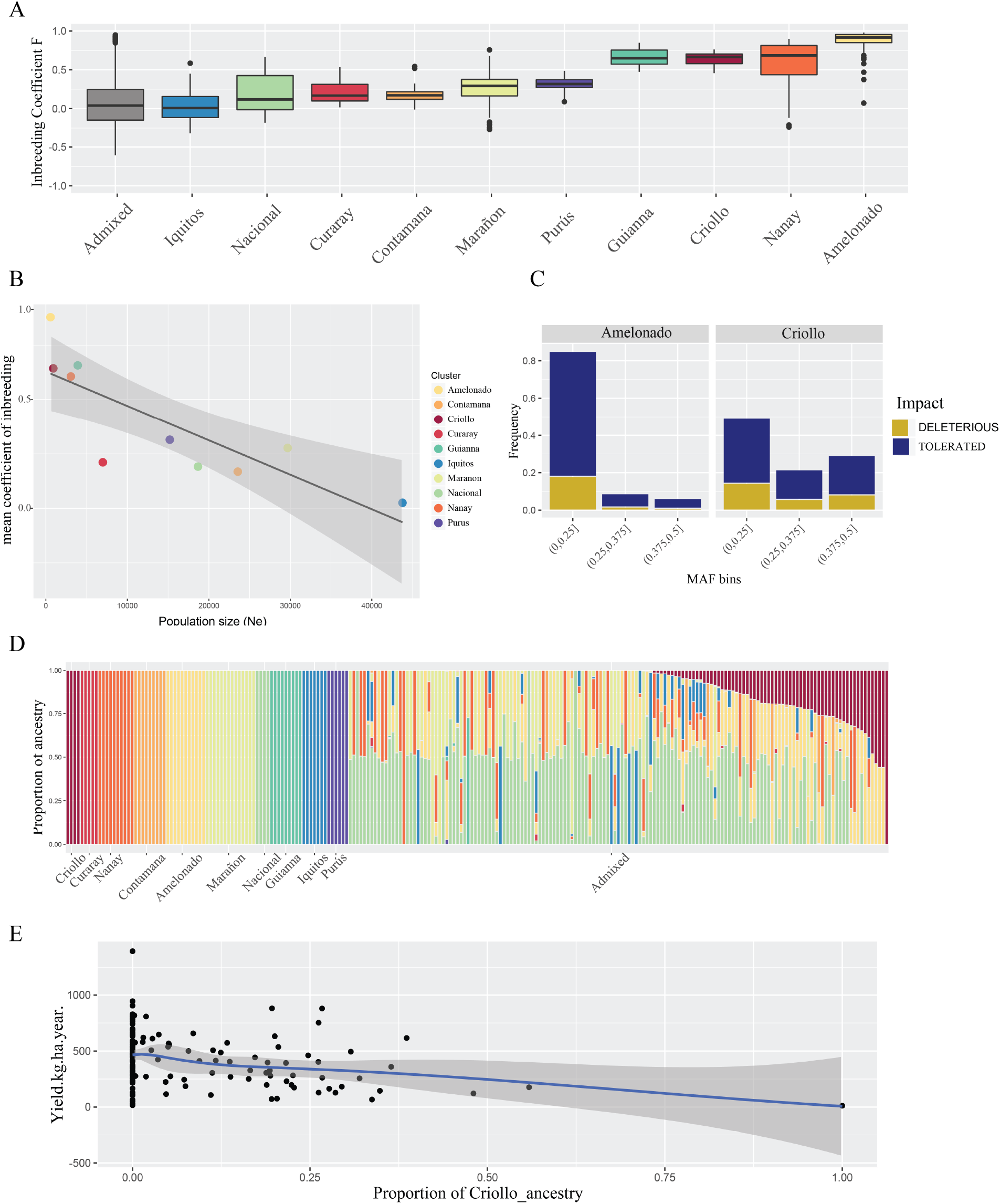
Accumulation of deleterious mutations during domestication in *T. cacao.* **A**, Distribution of coefficients of Inbreeding (F) per population (including the group of Admixed individuals). **B**, Coefficients of Inbreeding as a function of the harmonic mean of the effective population size (estimated from the median PSMC shown in Figure 2D). **C**, Distribution of deleterious/tolerated mutations inferred with SIFT for the Criollo and Amelonado groups for rare and two classes of common binned minor allele frequency classes showing the highest relative proportion of common deleterious and tolerated amino acid changes in Criollo. **D**, Population structure inferred using a maximum likelihood under a supervised model for a new set of genotyped individuals for which productivity has been measured. **E**, Productivity (measured as Kg of beans per hectare per year) as a function of Criollo ancestry in the newly genotyped set of individuals; the results show a significant reduction in productivity as the proportion of Criollo ancestry increases, after correcting for inbreeding.

Population genetics theory predicts that selfing increases the efficiency in the elimination of recessive deleterious mutations, when compared to outcrossing populations, because variants otherwise hidden in heterozygous individuals will be exposed to the action of natural selection^41, 42^. In contrast, domestication is a process that has been shown to contribute to the maintenance of deleterious mutations in higher frequency in populations^3, 39^. The impact of domestication on arboreal crops is not well understood, and it is even less understood in a plant like cacao that in domesticated varieties utilizes self-compatibility, a mechanism that tends to purge deleterious mutations. In order to test the impact of selfing and domestication on the accumulation of deleterious mutations in cacao, we annotated amino acid changes in *T. cacao* based on phylogenetic conservatism (as implemented in SIFT4, see methods) either as tolerated or deleterious (see Supplementary materials and methods).

We inferred the distribution of deleterious/tolerated changes for combined categories of minor allele frequency in Amelonado and Criollo. Amelonado was used to generate an expectation on the accumulation of deleterious mutations for a scenario with strong selfing (similar to Criollo) in the absence of strong domestication to understand the impact of domestication in the distribution of deleterious/tolerated in Criollos (Figure 4C). Amelonados present a distribution of deleterious/tolerated mutations with a high frequency of rare variants and a reduced representation of variants in intermediate and large minor allele frequencies. This is consistent with most deleterious mutations being purged by selfing in the population. On the other hand, we observed significant, larger counts of deleterious mutations in higher minor allele frequency classes in Criollo (Figure 4C and supplementary Figure S7), an observation that was significant across frequency classes (Mantel-Haenszel test, p-value < 2.2e-16, supplementary Figure S7). These differences indicate that selfing in the Criollo populations, despite being a predominant form of mating has not been a force strong enough to purge deleterious mutations in the population as was apparent in Amelonados. Similar patterns of accumulation of deleterious mutations have been reported in animals and plants, such as in the comparison of teosinte and domesticated maize^43^, but is reported here for the first time for an arboreal crop. In most of the analyses that have been performed to date in other organisms, including dogs and humans^39, 44^, it has not been shown what the impact of the accumulation of deleterious mutations is on fitness. We tested the hypothesis that the accumulation of deleterious mutations due to domestication would decrease fitness by examining the relationship between Criollo ancestry and a measure of performance in cacao using an independent dataset. We measured bean (seed) productivity (yield in kilograms of bean per hectare per year for each plant) as a measure of fitness. We inferred proportional ancestry to a new set of admixed individuals for which productivity had been assessed (Figure 4D), and show that there is a significant negative relationship between Criollo ancestry and fitness (Figure 4E, with criollo ancestry decreasing yield per hectare per year in ~319.9 units per percent unit of ancestry, p = 0.000425).

In summary, we provide the first general view of how natural diversification has shaped genetic variation in *Theobroma cacao.* We provide the first comprehensive view of the demographic scenario involved in the domestication of the Criollo variety, and identify genes that could function in the desirable taste attributes of the Criollo variety. We also pioneer, for arboreal crops, the use of genomic resources to evaluate how the process of domestication has shaped the pattern of accumulation of deleterious mutations, impacting fitness in a considerable way.

## METHODS SUMMARY

We sampled leaves from accessions at the Cacao Research Unit at the University of West Indies and CATIE in Costa Rica (Supplementary Information sections 1, 2). From DNA extracts we generated Illumina sequencing libraries, which were sequenced on the Illumina HiSeq platform (Supplementary Information sections 3, 4). We modified existing analysis pipelines for the identification of single nucleotide polymorphisms (Supplementary Information Section 5). To characterize genetic variation in the species we developed bash/R scripts and used existing tools for analyzing the population structure (Supplementary Information section 6). To test the association between genetic diversity and population differentiation we adjusted a generalized linear model (as described in Supplementary Information section 7). We investigated the evolutionary history of the populations using pairwise sequentially markovian coalescence (psmc) and adjusting a smoothing spline across individual histories inferred for each sample that corresponded to the same population (Supplementary Information section 8). We also use approximations based on maximum likelihood to investigate the evolutionary history of the differentiation between Criollo and Curaray to identify the most likely scenario that would explain the origin of the domesticated cacao (Supplementary Information section 9). We inferred regions under selection during domestication using existing methods that identify differences in the local pattern of genetic variation and the genome-wide pattern of genetic variation (Supplementary Information section 10). We used the estimated evolutionary history of the populations as inferred from psmc to estimate the harmonic mean of the effective population size for each genetic group in a comparable period of time, and adjusted a generalized linear model to explain the magnitude of inbreeding with difference in effective population size (Supplementary Information section 11). We inferred the annotation of deleterious and tolerated mutations in cacao after building a customized database for cacao using SIFT4 to analyze the differences in proportions of tolerated/deleterious mutations between Criollo and Amelonado (Supplementary Information section 12). The impact of the accumulation of deleterious mutations on productivity was assessed by fitting a generalized linear model to explain productivity (measured in Kg of beans/per hectare/per year) as a function of Criollo ancestry after correcting for inbreeding (Supplementary Information section 12). Statistical analyses were performed in R. SNP data will be made available in dbSNP format at the time of publication.

## Acknowledgements

All accessions of cacao sequenced in this work were obtained from the Cocoa Research Center (University of West Indies, Trinidad), the Centro Agronómico Tropical de Investigación y Educación (CATIE, Costa Rica) and samples obtained from NARS in Cameroon and the Ivory Coast. MARS provided with all the necessary funds to sequence accessions via a contract with CDB, and OEC was supported via this grant.

